# Highly-Neutralizing COVID-19-Convalescent-Plasmas Potently Block SARS-CoV-2 Replication and Pneumonia in Syrian Hamsters

**DOI:** 10.1101/2021.06.29.450453

**Authors:** Yuki Takamatsu, Masaki Imai, Kenji Maeda, Noriko Nakajima, Nobuyo Higashi-Kuwata, Kiyoko Iwatsuki-Horimoto, Mutsumi Ito, Maki Kiso, Tadashi Maemura, Yuichiro Takeda, Kazumi Omata, Tadaki Suzuki, Yoshihiro Kawaoka, Hiroaki Mitsuya

## Abstract

Despite various attempts to treat SARS-CoV-2-infected patients with COVID-19-convalescent plasmas, neither appropriate approach nor clinical utility has been established. We examined the efficacy of administration of highly-neutralizing COVID-19-convalescent plasma (*hn*-plasmas) and such plasma-derived IgG administration using the Syrian hamster COVID-19 model. Two *hn*-plasmas, which were in the best 1% of 340 neutralizing-activity-determined convalescent plasma samples, were intraperitoneally administered to SARS-CoV-2-infected hamsters, resulting in significant reduction of viral titers in lungs by up to 32-fold as compared to the viral titers in hamsters receiving control non-neutralizing plasma, while with two moderately neutralizing plasmas (*mn*-plasmas) administered, viral titer reduction was by up to 6-fold. IgG fractions purified from the two *hn*-plasmas also reduced viral titers in lungs than those from the two *mn*-plasmas. The severity of lung lesions seen in hamsters receiving *hn*-plasmas was minimal to moderate as assessed using micro-computerized tomography, which histological examination confirmed. Western blotting revealed that all four COVID-19-convalescent-plasmas variably contained antibodies against SARS-CoV-2 components including the receptor-binding domain and S1 domain. The present data strongly suggest that administering potent-neutralizing-activity-confirmed COVID-19-convalescent plasmas would be efficacious in treating patients with COVID-19.

**Importance:** Convalescent plasmas obtained from patients, who recovered from a specific infection, have been used as agents to treat other patients infected with the very pathogen. To treat using convalescent plasmas, despite that more than 10 randomized-controlled-clinical-trials have been conducted and more than 100 studies are currently ongoing, the effects of convalescent plasma against COVID-19 remained uncertain. On the other hand, certain COVID-19 vaccines have been shown to reduce the clinical COVID-19 onset by 94-95%, for which the elicited SARS-CoV-2-neutralizing antibodies are apparently directly responsible. Here, we demonstrate that highly-neutralizing-effect-confirmed convalescent plasmas significantly reduce the viral titers in the lung of SARS-CoV-2-infected Syrian hamsters and block the development of virally-induced lung lesions. The present data provide a proof-of-concept that the presence of highly-neutralizing antibody in COVID-19-convalescent plasmas is directly responsible for the reduction of viral replication and support the use of highly-neutralizing antibody-containing plasmas in COVID-19 therapy with convalescent plasmas.

## Introduction

More than a year had passed since the World Health Organization (WHO) declared a state of emergency, the pandemic of the novel coronavirus (severe acute respiratory syndrome coronavirus 2; SARS-CoV-2) disease (COVID-19) is still spreading worldwide^1,2^. More than 176 million people have been infected and more than 3.8 million lives have been lost by June 17, 2021 (https://covid19.who.int/) and COVID-19 is continuously posing most serious public health and socioeconomic problem globally in this century^3^. Vaccination is one of the most effective prophylactic health measures^4,5^ and considered as one of the most promising key strategy for curbing the current pandemic^6^. Multiple COVID-19 vaccines, such as mRNA vaccines, BNT162b2^7^, mRNA-1273^8^, ChAdOx1 nCoV-19/AZD1222^9^, and an adenovirus vector, Ad26.COV2.S^10^ are presently available in the US, the European Union, and other parts of the world. Different classes of vaccines such as a recombinant protein nanoparticle vaccine, NVX-CoV2373^11^, and inactivated COVID-19 vaccines, BBIBP-CorV^12^, CoronaVac^13^, and Covaxin^14^ are currently under development. Yet, how long the observed efficacy of vaccines lasts and whether such vaccines are effective in treating already-infected individuals remain to be determined^15^. In addition, the spread of SARS-CoV-2 variants which resist to the efficacy of certain vaccines throughout the world has been of great concern^16,17^.

Moreover, in terms of disease management, remdesivir^18^, dexamethasone^19^, baricitinib^20^, and IL-6 pathway inhibitors (e.g., tocilizumab)^21^ are the only recommended agents for severely ill patients with COVID-19, although the efficacy of such agents is only limited^22,23^ and no COVID-specific therapeutics are likely to be available in the immediate future. In this regard, immunotherapies for certain cancers and autoimmune disorders are relatively well established^24^; however, there are only a few immunotherapy for infectious diseases, which were shown to be efficacious. The efficacy of plasma infusions of SARS-CoV-1-convalescent plasma is controversial mainly because most clinical trials were not controlled or randomized^25^. Moreover, in many clinical trials, plasmas administered were not examined for their titers of neutralizing antibodies contained. Of note, fatality/clinical outcomes among those with COVID-19 receiving convalescent plasma whose titers of anti-SARS-CoV-2-receptor binding domain (RBD) antibodies have been reportedly lower than in those receiving no plasma^26,27^, especially when such plasmas were administered early after the onset. The outcomes of those receiving high-titer anti-SARS-CoV-2-RBD or anti-SARS-CoV-2-spike antibodies early after the onset have also been shown to be favorable^27^.

Imai and his colleagues have recently reported that SARS-CoV-2 efficiently replicates in the lungs of Syrian hamsters and causes severe pathological lung lesions that share the characteristics with lung lesions in patients with COVID-19^28^. Here, we examined the efficacy of neutralizing activity-confirmed COVID-19-convalescent plasmas and such plasma-derived IgG fractions by employing the SARS-CoV-2-exposed VeroE6 cells^29^ and Syrian hamster model. The present data strongly suggest that the treatment of COVID-19 patients using highly neutralizing activity-confirmed convalescent plasmas would efficiently block the development of COVID-19-associated lung lesions.

## Results

### COVID-19-convalescent-plasma-derived IgG fractions block SARS-CoV-2 infection in vitro

We have previously examined the presence and temporal changes of the neutralizing activity of IgG fractions from 43 COVID-19-convalescent plasma samples using cell-based assays^30^. In the current study, we chose two highly-neutralizing plasma (*hn*-plasma) samples and IgG fractions from Donor-043 (D43) and D84, which were in the best 1.4 and 0.5% of 340 neutralizing-activity-determined convalescent plasma samples, respectively, and two moderately-neutralizing plasma (*mn*-plasma) samples and IgG fractions from D73 and D91, which showed top 40.5 and 20.9% neutralizing activity in the 340 convalescent plasma samples, respectively, and confirmed their activity to block the cytopathic effect (CPE) of a SARS-CoV-2 strain (SARS-CoV-2^05-2N^) using VeroE6^TMPRSS2^ cells and the methyl thiazolyl tetrazolium (MTT) method^29,30^. Figure 1 shows that all the four representative COVID-19-convalescent plasmas and IgG samples significantly blocked the CPE of SARS-CoV-2^05-2N^. D43 and D84 plasmas were highly potent against the virus with IC_50_ values of 1,400±240 and 1,100±60 fold, respectively, while D73 and D91 samples showed relatively moderate activity with IC_50_ values of 220±30 and 400±90 fold, respectively (Figure 1a and Table 1). IgG fractions purified from D43 and D84 plasmas also exerted potent activity with IC_50_ values of 9.2±1.3 and 9.8±2.7 µg/ml, respectively, while those from D73 and D91 showed moderate activity with IC_50_ values of 47.9±9.0 and 24.9±3.1 µg/ml, respectively (Figure 1b and Table 1). A plasma sample from a healthy and qRNA-PCR-and-ELISA-confirmed SARS-CoV-2-uninfected individual and its IgG fraction failed to show significant CPE-blocking activity (Figure 1 and Table 1). We have also quantified the amounts of SARS-CoV-2-S1-binding antibodies in each plasma sample by using D84 plasma as a reference (100%) employing a commercially available ELISA kit. D43, D73, and D91 contained 140, 34, and 57% of IgG relative to D84 plasma (Table 1), showing that the amounts of S1-binding antibodies contained in plasma samples were roughly proportionate to the blocking effects of each plasma and IgG fraction, although it is of note that the presence of greater amounts of S1-binding antibodies in plasma does not necessarily predict the presence of greater levels of neutralizing activity^30^. Taken together, these data show that all the plasma samples used were highly or moderately active in blocking the infectivity and replication of SARS-CoV-2 and that the IgG fractions isolated from plasmas were largely responsible for the activity of plasmas to block the infectivity and CPE of the virus.

**Figure 1.**
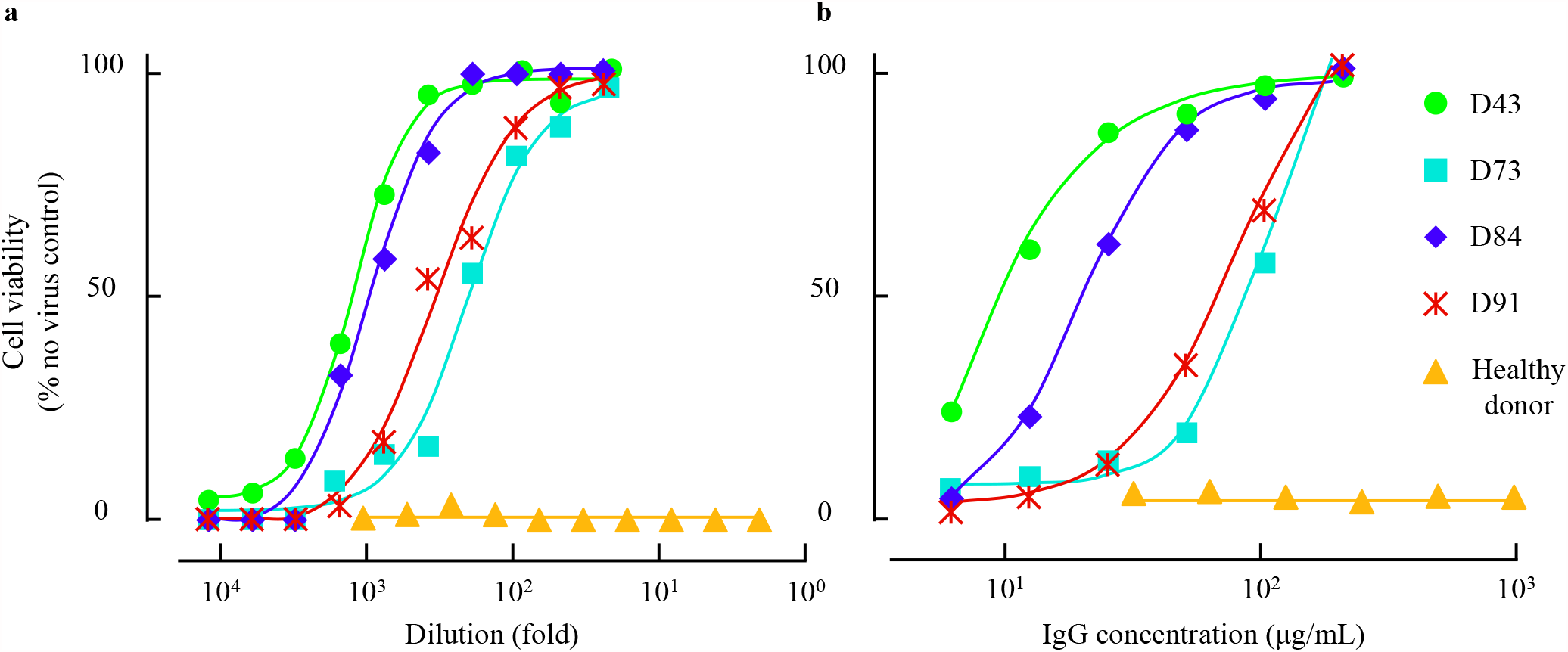
Anti-viral activity of convalescent plasma and purified IgG. VeroE6^TMPRSS2^ cells were exposed to SARS-CoV-2^05-2N^ with or without various concentrations of diluted plasma (**a**) or purified IgG (**b**). Note that highly neutralizing plasma (*hn*-plasma), D43 and D84, were highly potent while moderately neutralizing plasma (*mn*-plasma), D73 and D91, were relatively moderate active against the virus. A plasma sample from a healthy and qRNA-PCR-and-ELISA-confirmed SARS-CoV-2-uninfected individual and its IgG fraction failed to show significant CPE-blocking activity.

**Table 1.** The neutralizing activity of convalescent plasma and purified IgG. The neutralizing activity of COVID-19-convalescent plasmas and their IgG fractions were determined using MTT assay employing VeroE6^TMPRSS2^ cells. The relative amounts of anti-SARS-CoV-2-S1 binding antibody were quantified using anti-SARS-CoV-2-S1 IgG ELISA with serially diluted D84 plasma samples for standardization. All the convalescent plasma and purified IgG showed various potency of neutralizing activity, while healthy donor plasma and its purified IgG were inert against SARS-CoV-2^05-2N^. Of note, D43 derived plasma and purified IgG showed the most potent antiviral activity with the IC_50_ values of 1,400 ± 240-fold and 9.2 ± 1.3 μg/ml, respectively.

|  | Plasma IC<sub>50</sub><br>(fold) | Purified IgG IC<sub>50</sub><br>(µg/ml) | Anti-S1-IgG<br>(%) |
| --- | --- | --- | --- |
| D43 | 1,400 ± 240 | 9.2 ± 1.3 | 140 |
| D73 | 220 ± 30 | 47.9 ± 9.0 | 34 |
| D84 | 1,100 ± 60 | 9.8 ± 2.7 | 100 |
| D91 | 400 ± 90 | 24.9 ± 3.1 | 57 |
| Healthy donor | < 2 | > 1,000 | <0.3 |

### Body weight gains in SARS-CoV-2^UT-NCGM02^-exposed and neutralizing plasma-receiving Syrian hamsters were significantly greater than those in control-plasma-receiving animals

We have previously demonstrated that SARS-CoV-2 isolates efficiently replicate in the lungs of Syrian hamsters, causing severe pathological lung lesions^28^. Such SARS-CoV-2-infected 7- to 8-month-old hamsters also underwent substantial weight loss by day 7 post-infection and continued to lose weight for up to 14 days post-infection^28^. In the present study, we employed 1-month-old hamsters and intranasally inoculated them with 10^3^ plaque-forming units (PFU) of a clinically isolated SARS-CoV-2, SARS-CoV-2^UT-NCGM02^ (Set as Day 0). In 24 hours following the inoculation (on day 1), three hamsters per group were intraperitoneally administered with 2 ml of plasma from a qRNA-PCR-and-ELISA-confirmed SARS-CoV-2-uninfected healthy individual (control-plasma; See the protocol in Supp Figure 1). As the body weights of the control-plasma-receiving hamsters (n=3) were followed up every day, their weights continued to decrease by day 8 following the viral exposure, while the weights started to gain by day 9 and continued to gain thereafter. However, in hamsters that received the *hn*-plasma samples (D43 and D84), the decrease in body weights by day 8 was much less than in control-plasma-receiving hamsters and their body weights started to increase on day 9 and beyond (*p* values of the temporal changes in the body weights for the D43- and D84-receiving hamster groups to the control group were 0.0095 and 0.0092, respectively). One of the D73-plasma-receiving hamsters (Hamster#28) had a significantly greater body weight decrease among the four D73-plasma-receiving hamsters and the average body weights became close to those in the control-plasma-receiving hamsters (Figure 2; *p* value for the D73-plasma-receiving hamsters compared to four control-plasma-receiving hamsters was 0.2025).

**Figure 2.**
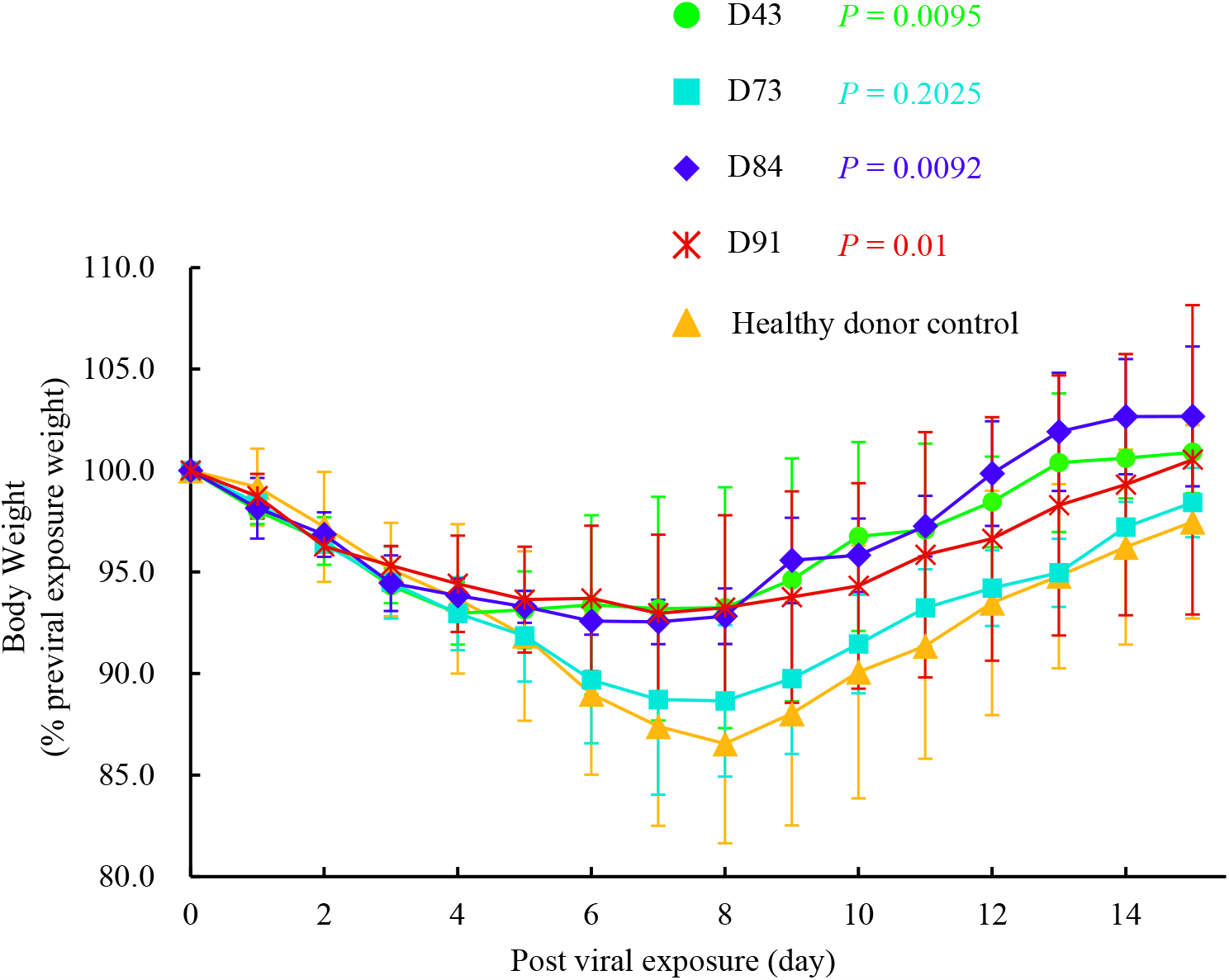
Body weight change in SARS-CoV-2 infected Syrian hamsters with plasma transfusion. Syrian hamsters were intranasally inoculated with 10^3^ PFU of clinically isolated SARS-CoV-2 (SARS-CoV-2^UT-NCGM02^). In 24 hours following the inoculation, hamsters were intraperitoneally administered with 2 ml of convalescent plasma or a qRNA-PCR-and-ELISA-confirmed SARS-CoV-2-uninfected healthy individual-derived control plasma, and the body weight was monitored daily for 15 days. The mean relative value from the pre-viral exposure baseline and S.D. values are shown. All the hamsters lost their weight by day 8 following the viral exposure, while the weights started to gain by day 9 and continued to gain thereafter. *P* values of the body weight change in D43-, D73-, D84-, and D91-receiving hamster groups to the control group were 0.0095, 0.2025, 0.0092, and 0.01, respectively.

### SARS-CoV-2^UT-NCGM02^-exposed and neutralizing plasma-receiving Syrian hamsters develop less severe pneumonia

SARS-CoV-2-infected Syrian hamsters undergo lung injuries, which share characteristics with injuries seen in the lungs of SARS-CoV-2-infected individuals, including severe, bilateral, largely peripherally distributed, multi-lobular ground glass opacity lesions and lobular consolidations as examined using microcomputed tomographic (micro-CT) imaging (Figure 3)^28^. In order to examine the effects of administering neutralizing human plasmas on the development of SARS-CoV-2-induced lung lesions in virus-exposed Syrian hamsters, we employed the *in vivo* X-ray micro-CT image capturing in the present study. In all three SARS-CoV-2^UT-NCGM02^-exposed hamsters, which intraperitoneally received 2 ml of the control-plasma on day 1 post-infection (Hamsters#21, #22, and #23), low-level infiltration with ground-glass opacities (GGOs) in bilateral lower lobes appeared by day 4 in both coronal and axial micro-CT thorax images (Supp Figure 2). By day 6, those lesions evolved into a mixed pattern of GGOs, consolidations, and interlobular septal thickening seen in whole lung. By day 8, such lesions further worsened to show GGOs with consolidations and fibrous stripes in bilateral lung accompanied with mediastinal emphysema, traction bronchiectasis, interlobular septal thickening, and/or cavitations (Supp Figure 2). Micro-CT scans taken on day 10, however, showed healing of the lung cavitation and mediastinal emphysema together with reduced GGOs. Micro-CT scans on day 12 show further healing of the consolidation and GGOs, while multiple focal fibrous stripes remained in bilateral peripheral field (Supp Figure 2).

**Figure 3.**
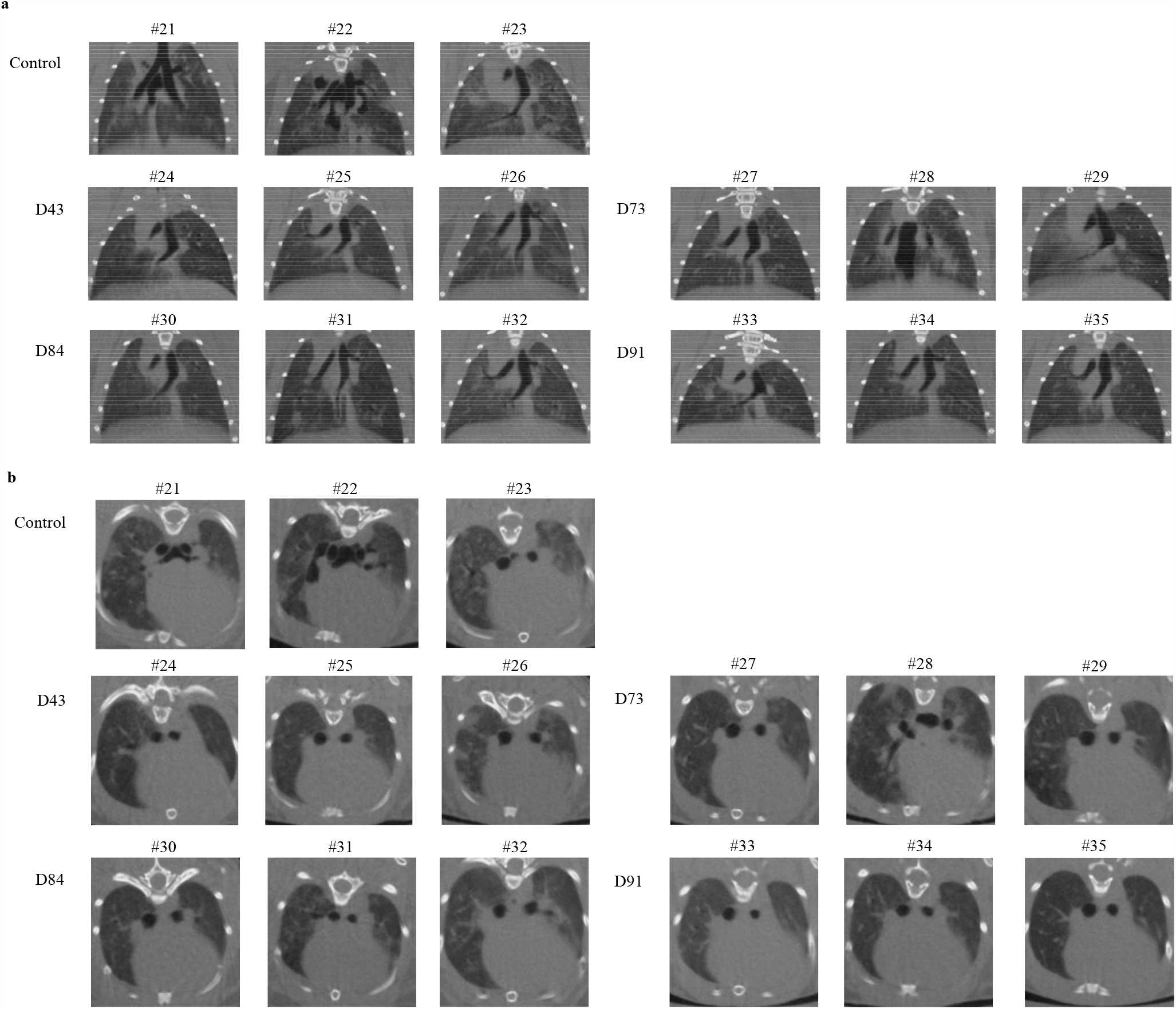
Micro-CT imaging of the lungs of SARS-CoV-2-infected Syrian hamsters with convalescent plasma transfusion on 8 days post viral exposure. (**a**) Coronal and (**b**) axial images of the thorax of hamsters receiving the SARS-CoV-2 inoculation and COVID-19-convalescent plasma i.p. transfusion. (**a, b**) The control plasma-receiving hamsters (Hamsters #21, #22, and #23) developed the ground-glass opacities (GGOs) with consolidations and fibrosis. However, in all three hamsters receiving *hn*-plasma (D43 for Hamsters #24, #25, and #26, and D84 for Hamsters #30, #31, and #32), no such lung abnormalities were observed throughout the 12 days micro-CT imaging in Hamster #24, #25, #30 and mild to moderate GGO and interlobular septal thickening were focally observed in #26, #31, #32. On the other hand, the chest CT images in *mn*-plasma receiving hamsters (D73 for Hamsters #27, #28, and #29, and D91 for Hamsters #33, #34, and #35) showed mixed but moderate GGO lesions and interlobular septal thickening in whole lung, however, no mediastinal emphysema or traction bronchiectasis were observed except in Hamster #28.

However, in all three hamsters that intraperitoneally received either of the two *hn*-plasma samples (Hamsters#24, #25, and #26 received 2 ml of D43 plasma; while Hamsters #30, #31 and #32 received 2 ml D84 plasma), no such extensive lung lesions developed throughout the 12-day period of observation and the difference between the lung images of D43- and D84-plasma-receiving hamsters and those of control-plasma-receiving Hamsters#21, #22, and #23 on day 8 post-infection (the dorsal lung images of control-plasma-receiving hamsters were taken from Supp Figure 2) was readily noticeable.

The lung CT scan images of *mn*-plasma-receiving hamsters (Hamsters#27, #28, and #29 received D73 plasma; while Hamsters #33, #34 and #35 received D91 plasma) showed mixed but moderate GGO lesions and interlobular septal thickening in whole lung, however, no mediastinal emphysema or traction bronchiectasis were observed except in Hamster#28 (Figure 3a). Coronal micro-CT scan images confirmed the moderate changes in the lung scan images of those hamsters as compared to the lung CT images of the control-plasma-receiving hamsters (Figure 3b).

### D43-plasma administration apparently inhibited the spread of viral infection from bronchiolar to alveolar regions

On day 4 post-infection (on day 3 post-plasma-administration), histopathological features and virus distribution pattern in the lung tissues of each hamster were examined. All histopathology and immunohistochemistry features obtained are illustrated in Supp Figure 3. Histopathology of the lung sections of each animal showed moderate inflammatory cell infiltration consisting of neutrophil, monocytes/macrophages, and lymphocytes around the bronchi and bronchioles. In some regions, the inflammatory cells were detected in the alveoli. However, the degrees of histopathological changes substantially varied among hamsters, and there was no readily significant difference among the groups administered with different plasmas (Figure 4, panels a, c, e, g, i, and Supp Figures 3a–e). Then, immunohistochemistry with anti-SARS-CoV-2 antibody revealed viral antigens in the bronchiole epithelium and viral spreading to the alveolar epithelium surrounding the bronchioles in all the animals except for D43-plasma-receiving hamsters. Notably, in the D43-receiving animals (Figure 4d and Supp Figure 3b), substantially less viral antigens were seen and the extent of the viral spreading was apparently limited to bronchial and alveolar epitheliums adjacent to bronchioles regardless of the degrees of histopathological changes compared to the amounts of viral antigens seen in the lungs of control-plasma-, and D73-, D84-, and D91-plasma-receiving hamsters (Figure 4b, f, h, j, and Supp Figures 3a, c, and d). Moreover, the numbers of viral antigen-positive-cells in the alveolar regions also appeared less in the lung of D43-plasma-receiving hamsters (Figure 4d and Supp Figure 3b) than in the control-plasma-receiving and D73-, D84-, or D91-plasma-receiving animals (Figure 4b, f, h, j, and Supp Figures 3a, c, and d).

**Figure 4.**
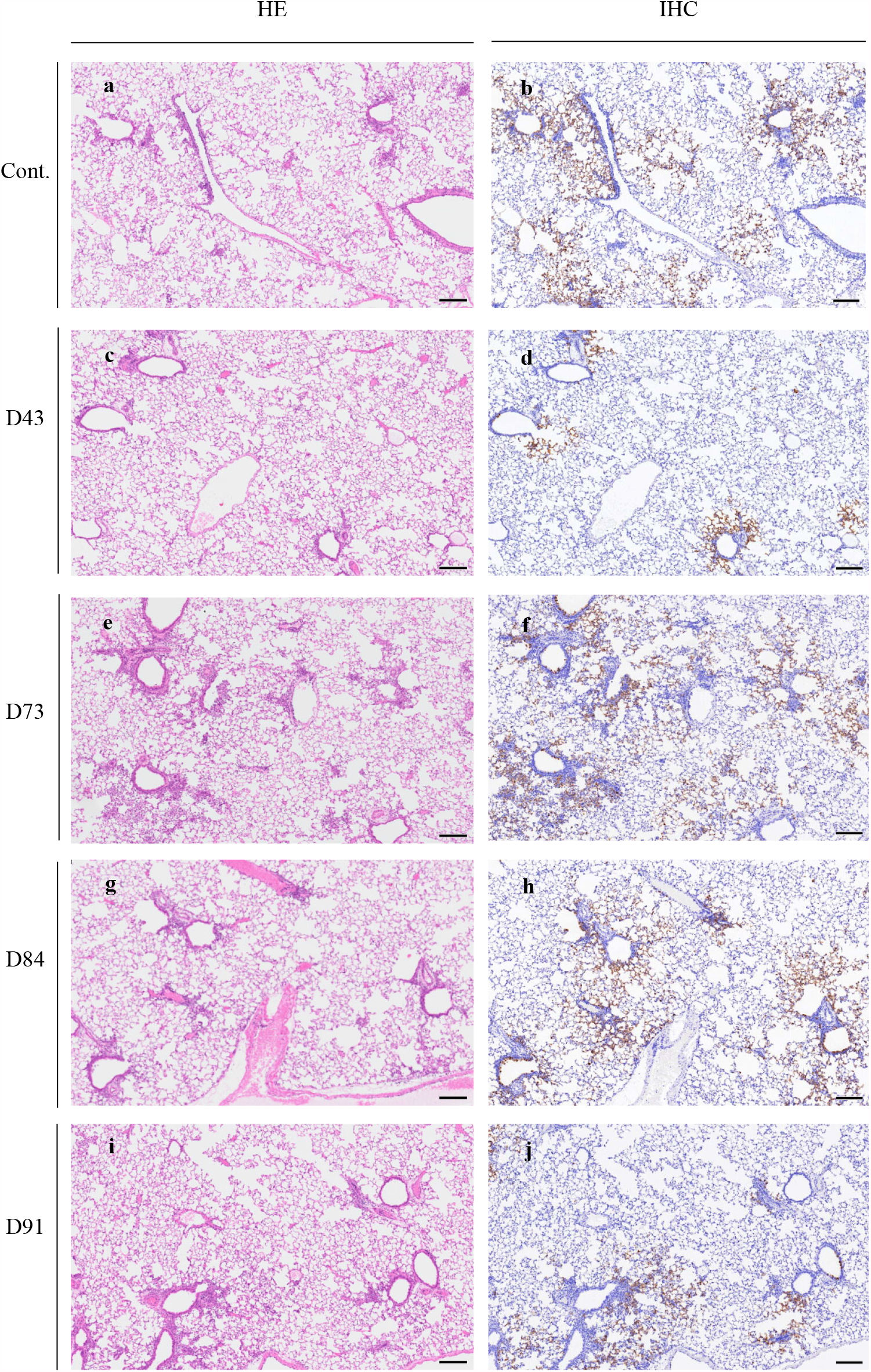
Pathological examination of the hamsters treated with plasma. Representative images of histopathology and immunohistochemistry on the lung sections of hamsters treated with control or each plasma on Day 1 post-infection are shown. Hematoxylin Eosin (HE) staining of the lung sections obtained from the control (**a**) and plasma D43 (**c**), D73 (**e**), D84 (**g**), or D91 (**i**) treated animals. Immunohistochemistry (IHC) for SARS-CoV-2 antigen detection of the lung sections obtained from the control (**b**) and plasma D43 (**d**), D73 (**f**), D84 (**h**), or D91 (**j**) treated animals. Scale bar = 200μm.

### Neutralizing activity-confirmed plasmas significantly suppressed the replication of SARS-CoV-2 in the lung of hamsters

All the neutralizing activity-confirmed COVID-19-convalescent plasma samples (D43, D73, D84, and D91 plasmas) mitigated the body weight reduction and SARS-CoV-2-induced lung lesions (Figure 2, Supp Figure 2, and Figure 3); however, the histopathological examination and immunostaining method largely failed to detect differences in the presence or spread of the virus between the hamsters receiving the control-plasma and those receiving neutralizing-activity-confirmed plasmas (Figure 4 and Supp Figure 3). Thus, we attempted to quantify the amounts of infectious virions in the lungs of hamsters receiving control-plasma, D43-, D73-, D84-, or D91-plasma samples. Each hamster was exposed to the virus on day 0, intraperitoneally administered with 2 ml of each plasma on day 1, and sacrificed on day 4. Thereafter, each lung was homogenized and the virus titers in the homogenates were determined employing plaque forming assays using VeroE6^TMPRSS2^ cells. As shown in Figure 5a, the geometric mean titer for the hamsters receiving control-plasma was 10^8.5^ PFU/g, while the administration of D43 and D91 plasmas had significantly suppressed the replication of SARS-CoV-2^UT-NCGM02^ with viral titers of down to 10^7.0^ (*p*=0.0003) and 10^7.7^ (*p*=0.037) PFU/g, respectively, while the reductions by D73 and D84 plasmas were not statistically significant (*p*>0.05; Figure 5a). When IgG fractions isolated from plasmas were intraperitoneally administered to hamsters, D43 IgG fraction gave the greatest reduction with a geometric mean infectious viral titer of 10^7.1^ PFU/g compared to the viral titer in the control-plasma-receiving hamsters with a geometric mean titer of 10^8.4^ PFU/g; while D91, D84, and D73 IgG fractions gave mean titers of 10^7.7^ (*p*=0.015), 10^7.8^ (*p*=0.037), and 10^8.0^ (*p*>0.05) PFU/g, respectively (Figure 5b).

**Figure 5.**
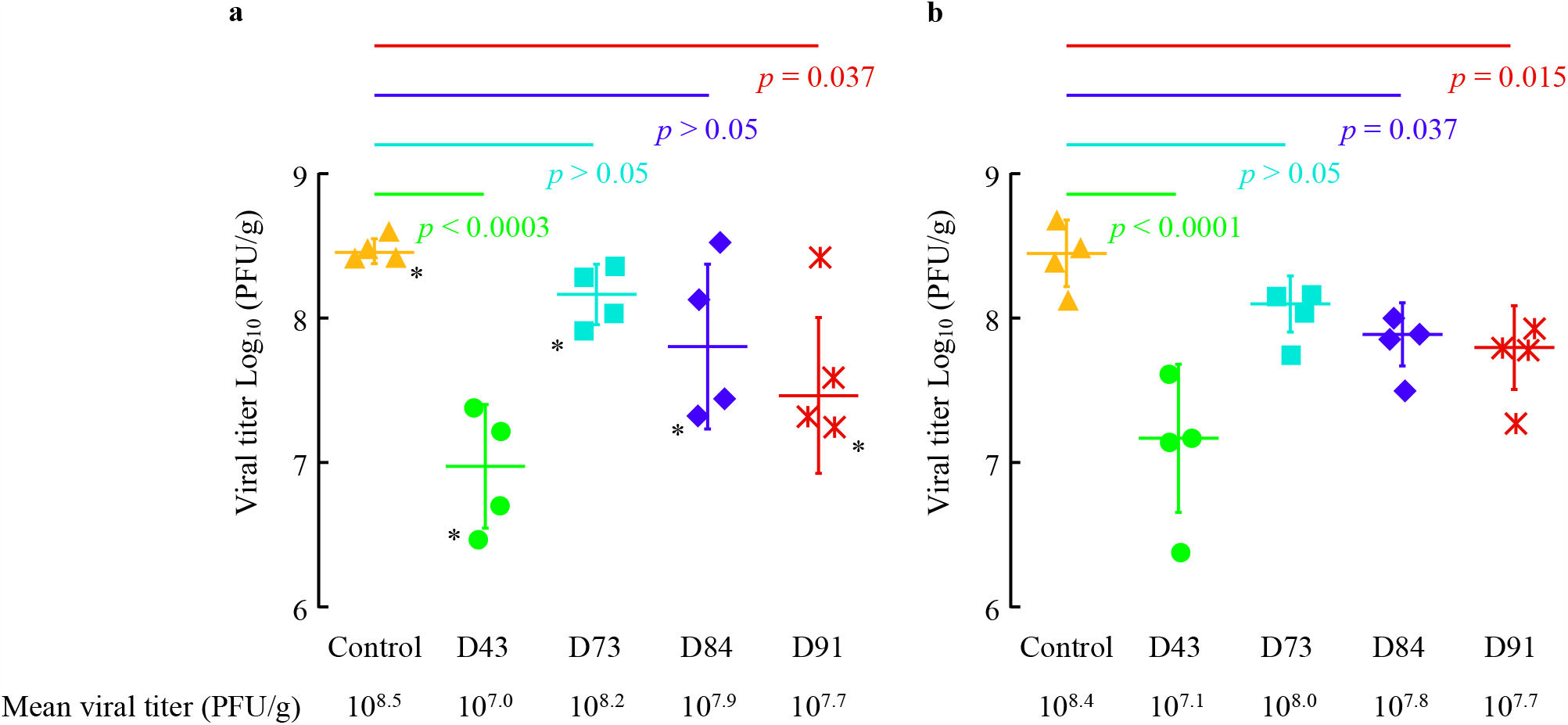
Neutralizing activity of convalescent plasma in the lungs of SARS-CoV-2 infected Syrian hamsters with plasma transfusion. Syrian hamsters were intranasally inoculated with 10^3^ PFU of clinically isolated SARS-CoV-2 (SARS-CoV-2^UT-NCGM02^). In 24 hours following the inoculation, hamsters were intraperitoneally administered with 2 ml of plasma (**a**) or plasma-derived purified IgG (**b**). Four Syrian hamsters per group were sacrificed on 4 days post viral exposure (3 days post plasma transfusion), and the virus titers in the lungs and neutralizing antibody titer in sera were determined by employing VeroE6^TMPRSS2^ cells. The geometric mean titer and S.D. values are shown.

In an attempt to see the effects of administering IgG fractions isolated from neutralizing human plasmas on the development of lung lesions in virus-exposed animals, we conducted additional histopathological and immunostaining study. Since we had failed identifying the difference in the histopathological findings among lungs of hamsters receiving various plasmas, we examined the lung in one hamster, which had the lowest infectious viral titer among each group (n=4) in this additional study. Representative images of the immunostained lung sections of hamsters showed that the infected cells are observed from the terminal bronchioles into the alveolar region in animals treated with control-plasma IgG, IgG from D73, D84, and D91 plasmas; however, the number of infected cells was much less in the terminal bronchioles and alveolar regions in the hamster receiving IgG fraction from D43 plasma (Supp Fig 4), corroborating the histopathological and immunostaining observations in hamsters receiving plasmas (Figure 4).

### COVID-19-convalescent-plasmas variably contain antibodies that specifically bind to viral components

We finally attempted to determine which antibodies within the four convalescent D43, D73, D84, and D91 plasma samples bind to SARS-CoV-2 components using the Jess capillary-based Western blot system. The four viral components (RBD, S1, S2, and nucleocapsid) are covalently fixed to the capillary and the presence of human IgG specifically bound to each viral component in the capillary is detected by exposing the capillary to HRP-conjugated anti-human IgG and iridescent light elicited by luminol being mediated by HRP. Figure 6a illustrates that each of the plasma contained IgG antibodies reactive with viral components, showing that the amounts and the ratios of each viral component-specific antibodies were substantially varied. Among the four convalescent plasma tested, D43, one of the two *hn*-plasmas contained the highest amounts of anti-RBD, -S1, and -NC IgG, while D84 contained the highest amount of anti-S2 and -whole Spike IgG (Figure 6b). Interestingly, the two *mn*-plasmas contained low levels of anti-RBD, -S1, -whole spike and -NC IgG.

**Figure 6.**
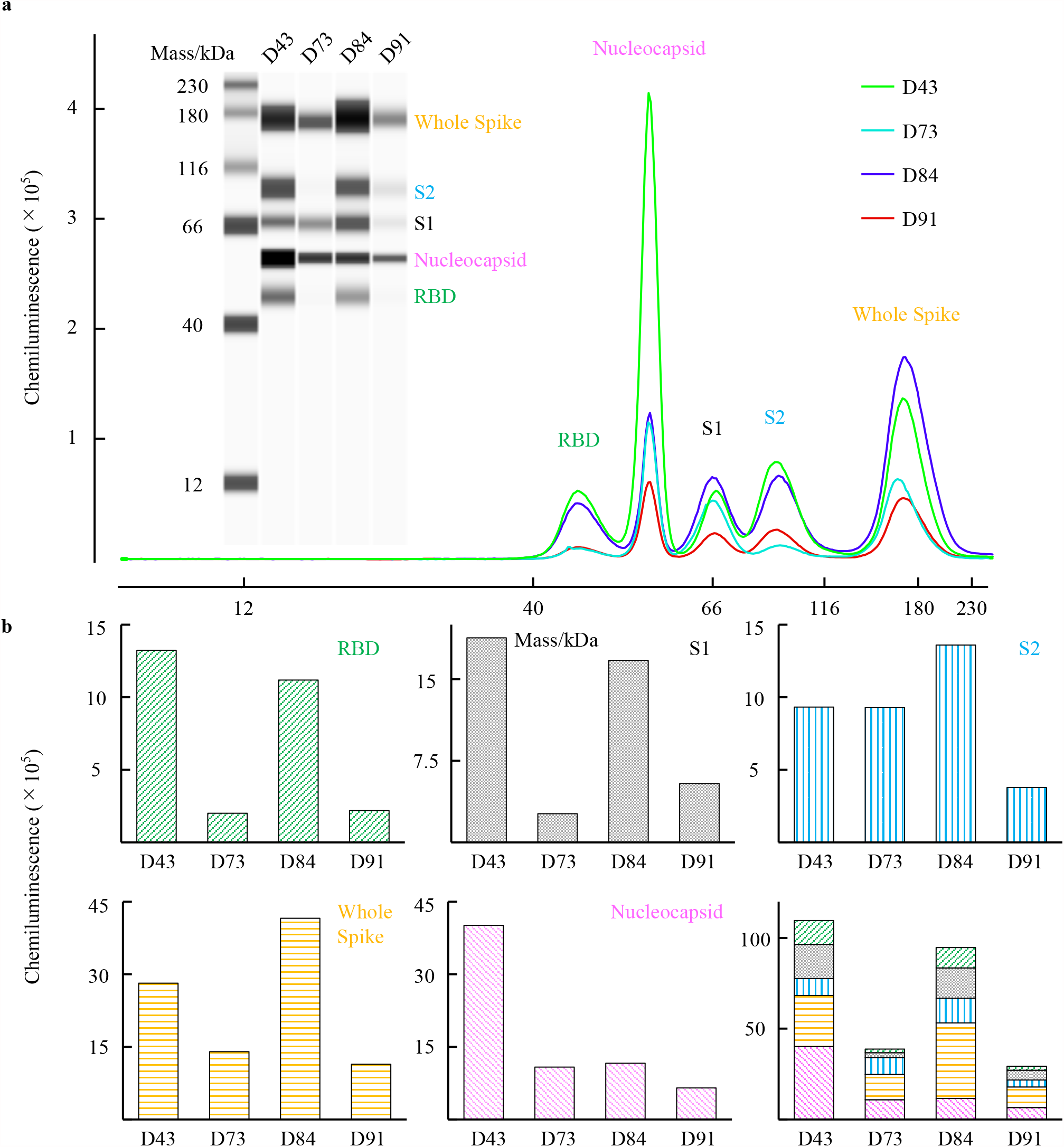
The characteristics of the anti-SARS-CoV-2 IgG in convalescent plasma against multiple viral components. The anti-SARS-CoV-2 IgG in each COVID-19-convalescent plasma reactive against SARS-CoV-2 viral components (RBD, S1, S2, whole Spike, and nucleocapsid [NC]) was detected using the Simple Western Jess System. (**a**) Western blotting image obtained with the Jess System and the immunoreactive signals for the presence of viral components bound by anti-SARS-CoV-2 IgG by using Simple Western Jess System. All four COVID-19-convalescent plasmas contained the SARS-CoV-2-specific IgG. (**b**) The quantification of IgG levels in four COVID-19-convalescent plasmas. D43 and D84 plasma samples had high amounts of anti-RBD IgG, while D73 and D91 had much less amounts of anti-RBD IgG. The same trend was seen in the amounts of anti-S1 and anti-whole Spike IgGs. On the other hand, D43 sample had lower amount of anti-S2 IgG than D84. D43 had a much higher amount of anti-NC IgG compared with D73, D84, and D91.

## DISCUSSION

In the present study, we demonstrate that highly neutralizing COVID-19-convalescent plasmas (*hn*-plasmas) reduced the severity of lung lesions in SARS-CoV-2-exposed Syrian hamsters compared to those receiving a non-neutralizing (control) plasma or moderately-neutralizing plasmas (*mn*-plasmas) as assessed with microCT-captured images (Supp Figure 2 and Figure 3) and the presence of SARS-CoV-2-infected cells in the lung (Figure 4 and Supp Figures 3 and 4). Moreover, *hn*-plasmas induced significant reduction of viral titers in the lungs of SARS-CoV-2-exposed Syrian hamsters as compared to those receiving control plasma or *mn*-plasmas (Figure 5a). IgG fractions purified from *hn*-plasmas also substantially reduced viral titers in the lungs of hamsters (Figure 5b). These data strongly suggest that administering *hn*-COVID-19-convalescent plasmas would be efficacious in treating patients with COVID-19 and mn-plasmas are unlikely to be effective in treating COVID-19 patients. The data also suggest that the IgG fractions largely contribute to the antiviral activity of *hn*-plasma, although other immune responses such as CD8^+^ killer T-cell and Fc-effector functions may contribute to protection and their relative importance in protection against COVID-19 is to be investigated^33,34^.

Several recent clinical studies suggest that neutralizing antibodies are generally sufficient to confer protection against the SARS-CoV-2 infection and that the protection against COVID-19 development is largely explained by SARS-CoV-2-neutralizing antibody responses, leaving less room for impact of T cells on correlation.

Yet, the efficacy of infusions of COVID-19-convalescent plasma has been controversial mainly because most clinical trials were not well controlled or randomized^25^. However, the failure of most COVID-19-convalescent plasma infusion studies to prove to be efficacious is likely due to the facts that the plasmas used were not confirmed to contain high titers of neutralizing activity before transfusion. In fact, in a few studies, in which only convalescent plasma whose levels of anti-SARS-CoV-2-RBD antibodies were confirmed to be high, have produced favorable clinical results^26,27^.

It is noteworthy that even in the well-planned clinical studies where high titers of neutralizing antibodies were used, a number of such clinical studies employed different ways and means to express neutralizing activity such as 50% neutralization titers, reciprocal neutralizing antibody titers, and IC_50_ log_10_ geometric mean titers. Moreover, such clinical studies have used different cells and viral strains in quantifying neutralizing activity of plasmas. Thus, establishing “international unit” or “international standard” is being thought to be needed to compare across various studies and to calibrate the strength of neutralization with a reference human convalescent sera panel. Establishing such standard should improve the correlation between the levels of neutralization activity and resulting clinical efficacy^35^.

In our previous study^30^, to possibly calibrate the neutralizing activity of COVID-19-convalescent plasmas with the neutralizing activity of plasmas determined in other studies, we used neutralizing unit per mg protein of IgG derived from such COVID-19-convalescent plasmas. In the present study, from among 340 COVID-19-convalescent plasma samples we have examined for their neutralizing activity using SARS-CoV-2^05-2N^ that was isolated in Tokyo in March 2020 and TMPRSS2-overexpressing VeroE6 (VeroE6^TMPRSS2^) cells as target cells as previously described^30^, we chose four plasma samples as references of SARS-CoV-2-neutralizing plasmas. The four plasma samples, D43, D73, D84, and D91 were in the top 1.4%, 40.5%, 0.5%, and 20.9% of 340 COVID-19-convalescent plasma samples, respectively. It is of note that approximately 60% of convalescent plasmas have low or no significant neutralizing activity as we mentioned in our previous report^30^. The reason why D43 was most effectively blocked the infection and replication of the virus in hamsters, while the comparably neutralizing D84 was less effectively blocked the virus in hamsters, remains to be elucidated. One possibility is that convalescent plasmas contain polyclonal neutralizing antibodies and their constituents may substantially vary from one convalescent patient to another^36,37^. Thus, even if a plasma sample exerts potent neutralizing activity in a cell-based assay, its neutralizing activity may not be directly well reproduced in the bodies of hamsters where the virus may unevenly infect and replicate so that the efficacy may vary depending on the constituent of polyclonal neutralizing antibodies.

One limitation of the current study is that only one SARS-CoV-2 strain (SARS-CoV-2^05-2N^) was employed and the results obtained here may not predict the efficacy in individuals infected with other SARS-CoV-2 strains, in particular, SARS-CoV-2 variants recently isolated^38–41^, which may escape neutralizing antibodies in plasmas used in the present study^42^ or may replicate more efficiently than previously isolated SARS-CoV-2 strains such as SARS-CoV-2^05-2N 43^. However, if convalescent plasmas are collected from individuals who are infected with certain SARS-CoV-2 variants, the plasmas from such individuals should be of use to immediately treat others infected with the same variants.

In conclusion, the present data strongly suggest that administering highly-neutralizing COVID-19-convalescent plasmas should be efficacious in treating patients with COVID-19, but potent neutralizing activity has to be confirmed before administering such convalescent plasmas.

## MATERIALS AND METHODS

### Patients

Four patients, who were clinically diagnosed with COVID-19 and agreed to participate in the clinical studies (approval number NCGM-G-003472 and NCGM-G-003536) for convalescent plasma donation, were selected^30^. Donated plasma was stored at -20°C until use.

### Cells, viruses, and IgG purification

TMPRSS2-overexpressing VeroE6 (VeroE6^TMPRSS2^) cells (RRID: CVCL_YQ49) were obtained from the Japanese Collection of Research Bioresources (JCRB) Cell Bank (Osaka, Japan). VeroE6^TMPRSS2^ cells were maintained in Dulbecco’s modified Eagle’s medium (DMEM) supplemented with 10% fetal bovine serum, 100 μg/ml penicillin, 100 μg/ml kanamycin, and 1 mg/ml G418 under humidified atmosphere containing 5% CO_2_ at 37°C. Two SARS-CoV-2 strains, hCoV-19/Japan/UT-NCGM02/2020 (SARS-CoV-2^UT-NCGM02^, GISAID Accession ID; EPI_ISL_418809)^28^ and 05-2N (SARS-CoV-2^05-2N^)^30^ were clinically isolated as previously described. IgG fractions were purified from convalescent plasma at Immuno-Biological Laboratories (Gunma, Japan) by using rProtein A Sepharose Fast Flow (Cytiva, Marlborough, MA) and eluted in phosphate-buffered saline (PBS). The IgG fractions were stored at -80°C until use.

### Antiviral assays

The SARS-CoV-2 neutralizing activity of donated plasma and purified IgG was determined as previously described^29–31^. In brief, VeroE6^TMPRSS2^ cells were seeded in 96-well flat microtiter culture plates at the density of 1 × 10^4^ cells/well. On the following day, the virus (SARS-CoV-2^05-2N^) was mixed to the various concentrations of the plasma or purified IgG fractions and incubated for 20 min at 37°C. The preincubated mixture was inoculated to the cells at a multiplicity of infection (MOI) of 0.01. The cells were cultured for 3 days and the number of viable cells in each well was measured using Cell Counting Kit-8 (Dojindo, Kumamoto, Japan). The potency of SARS-CoV-2 inhibition by plasma or purified IgG was determined based on its inhibitory effect on virally induced cytopathicity in VeroE6^TMPRSS2^ cells. The amounts of SARS-CoV-2-S1-binding antibodies in each plasma sample were determined by using Anti-SARS-CoV-2 ELISA (IgG) (Euroimmun, Lübeck, Germany). The total human IgG concentration was determined by using Human IgG ELISA Kit (abcam, Cambridge, UK).

### Experimental Infection of Syrian Hamsters

All the animal infection experiments were conducted as previously described^28^. In brief, one-year-old male Syrian hamsters (Japan SLC Inc., Shizuoka, Japan) were enrolled. Hamsters were intranasally inoculated with 10^3^ PFU (in 100 μl) of SARS-CoV-2^UT-NCGM02^ under ketamine-xylazine anesthesia. On the following day, 2 ml of convalescent plasma (experiments 1 and 2) or purified IgG (experiment 3) were intraperitoneally (i.p.) transfused to each Syrian hamster (Supp. Figure 1). The total dosage and the dosage per body weight of human IgG in 2 ml plasma are shown in Supp. Table 1 (Supp. Table 1). The total amount of human IgG in purified IgG fraction transfused to hamsters and plasma equivalent are shown in Supp. Table 2 (Supp. Table 2). The hamsters were monitored until the designated endpoint of the experiments.

For experiment 1, to monitor the body weight change and the micro-CT image, three hamsters per group were enrolled. The daily body weight was monitored for 15 days, and micro-CT imaging was conducted on days 0, 4, 6, 8, 10, and 12 post infection. The body weight was compared with the pre-infection baseline, and the relative values were calculated. The change in the body weights from the baseline of each hamster treated with plasma were compared and *p* value was calculated. In experiments 2 (plasma administered i.p.) and 3 (purified IgG fraction administered i.p.), in order to determine the *in vivo* antiviral activity of the convalescent plasma or purified IgG, four hamsters per group were enrolled. Hamsters were sacrificed on the fourth day post infection, and lungs were collected for histological examination and viral titration (Supp. Figure 1). The viral titer in the lungs was determined by means of plaque assays in VeroE6^TMPRSS2^ cells. All experiments with hamsters were performed in accordance with the Science Council of Japan’s Guidelines for Proper Conduct of Animal Experiments. The protocol was approved by the Animal Experiment Committee of the Institute of Medical Science, the University of Tokyo (approval number PA19-75).

### Micro-CT imaging

The chest CT images of the SARS-CoV-2-infected hamsters were captured as previously described using an *in vivo* micro-CT scanner (CosmoScan FX; Rigaku Corporation, Japan) until 12 days post-infection under ketamine-xylazine anesthesia. The imaging was conducted at the following conditions; 2 min at 90 kV, 88 μA, FOV 45 mm, and pixel size 90.0 μm. After scanning, the lung images were reconstructed by using the CosmoScan Database software (Rigaku Corporation) and analyzed as manufacturer’s instruction.

### Pathological examination

Excised animal tissues were fixed in 10%-buffered formalin and processed for paraffin embedding. The paraffin blocks were cut into 3-µm-thick sections and then mounted on silane-coated glass slides. One section from each tissue sample was stained using a standard hematoxylin and eosin procedure; another was processed for immunohistochemistry. After deparaffinization, antigens were activated (121°C, 10 min) with Target Retrieval Solution pH6.0 (Dako Cytomation, Glostrup, Denmark), and endogenous HRP was inactivated by hydroperoxide treatment. The sections were treated with 5 % normal goat serum for 30 minutes at room temperature and incubated with rabbit monoclonal anti SARS-CoV nucleoprotein antibody (Sino Biological, Beijing, China) at 4°C overnight. Specific antigen-antibody reactions were visualized by means of 3,3’-diaminobenzidine tetrahydrochloride staining using the Dako Envision system (Dako Cytomation).

### Detection and quantification of anti-SARS-CoV-2 IgG bound to viral components

The amounts of anti-SARS-CoV-2 IgG antibodies reactive with SARS-CoV-2 viral components in convalescent plasma were determined using the Simple Western Jess apparatus and the SARS-CoV-2 Multi-Antigen Serology Module (Protein Simple, San Jose, CA) according to the manufacturer’s instructions. In brief, various recombinant viral components [RBD (200 µg/ml), nucleocapsid [NC](5 µg/ml), S1(20 µg/ml), S2(20 µg/ml), and whole Spike (20 µg/ml)] were covalently fixed with ultraviolet irradiation to a 12-230 kDa Jess & Wes Separation Module (Protein Simple). The immobilized viral components were then exposed to each of 30-fold-diluted convalescent plasma samples (primary antibodies). Subsequently, the antibodies bound to the viral components were probed with horseradish peroxidase (HRP)-conjugated anti-human IgG (secondary antibody). The presence of human IgG in the Module is detected by iridescent light produced by luminol reagent being mediated by HRP. The quantification of each signal was performed using Compass for SW software ver. 5.0.1 (Protein Simple).

### Statistical analysis

For the comparison of the temporal changes in body weights of hamsters receiving control and convalescent plasma (control, D43, D73, D84, and D91), the changes in the body weights relative to the body weight before viral exposure were modeled with quartic functions. This is because each curve has a minimum around the middle of the post infection days and high values at its both edges. Therefore, the number of parameters determined is five, and the function was fitted to the data by use of the nonlinear least squares method which was performed by the Levenberg-Marquardt algorithm. The F statistics for the comparisons of two curves of the body-weight changes were calculated and the *p* values were derived^32^. The distribution of the residuals was tested and found to be consistent with normality. Because of the nonlinearity of the model, the *p* values are only approximate. For the viral titer in lung, each the convalescent plasma receiving group was compared with the healthy donor plasma receiving group using Dunnett’s test by using JMP Pro 15.0.0 (SAS Institute).

## Contributors

Conceptualization, Yuk.T, K.M., Y.K., and H.M.; Methodology, Yuk.T., M.I., K.M., N.N., Y.K., and H.M.; Formal Analysis, K.O.; Investigation, Yuk.T., M.I., K.M., N.N., N.H-K., K.I-H., M.I., M.K., T.M., and Yui.T.; Data curation, Yuk.T., M.I., K.M., K.I-H, M.I., T.M.; Writing – Original Draft, Yuk.T. and H.M.; Writing-Review & Editing, M.I., N.N., N.H-K., Yui.T., K.O., T.S., and Y.K.; Supervision, T.S., Y.K., and H.M.; Project Administration, H.M.; Funding Acquisition, K.M., Y.K. and H.M.

## Declaration of Interests

All authors declare that they do not have any competing interests related to this study.

## Acknowledgments

This work was supported in part by Japan Agency for Medical Research and Development (AMED) (grant numbers 20fk0108160 and 20fk0108502 to K.M.; JP19fk0108113, JP20nk0101612, JP19fm0108006, JP21wm0125002, JP20fk0108260 and JP20fk0108502 to Y.K.; and 20fk0108502, 20fk0108257, and 20fk0108510 to H.M.); by MHLW Research on Emerging and Re-emerging Infectious Diseases and Immunization Program (grant number JPMH20HA1006 to K.M.); by a grant from National Center for Global Health and Medicine Research Institute (grant number 20A2003D to K.M.) ; by National Institutes of Allergy and Infectious Diseases (grant number HHSN272201400008C to Y.K.). These funding sources were not involved in study design, in the collection, analysis, and interpretation of data, in the writing of the report, and in the decision to submit the paper for publication. We are grateful to Dr. Miwa Tamura-Nakano and Ms. Chinatsu Oyama in the communal laboratory of NCGM Research Institute for their technical support. The authors also thank Ms. Mariko Kato for technical assistance.

## Supporting Figure Legends

**Supp. Figure 1. Scheme of the Syrian Hamster experiments**. Hamsters were intranasally inoculated with 10^3^ PFU (in 100 μl) of SARS-CoV-2^UT-NCGM02^ (Set as Day 0). In 24 hours, 2 ml of convalescent plasma (experiments 1 and 2) or purified IgG (experiment 3) was intraperitoneally (i.p.) transfused to each Syrian hamster. In experiment 1, micro-CT imaging and the body weight monitoring were conducted for 15 days. In experiments 2 and 3, hamsters were sacrificed on day 4 and the histological examination and viral titration of lungs were conducted. The distribution of anti-viral activity of the 340 donated convalescent plasma samples and four convalescent plasma tested are indicated in violin plot.

**Supp. Figure 2. Administration of IgG fraction purified from plasma D43 effectively blocks the replication and spread of SARS-CoV-2 infection in lung tissue**. Fixed/paraffin-embedded lung tissue sections were immuno-histologically stained with anti-SARS-CoV-2-nucleoprotein polyclonal antibodies (in brown) and examined under light microscopy. Nuclei were counterstained with Mayer’s hematoxylin (in blue). Representative images of the immune-stained lung sections of hamsters receiving IgG fraction isolated from the control-plasma (a) and IgG fractions from D43 (b), D73 (c), D84 (d), and D91 (e) plasmas are shown. Each inset shows a higher magnification field of the rectangle area, showing the terminal bronchioles open into the alveolar region. Note that the infected cells are observed from the terminal bronchioles into the alveolar region in animals treated with control-plasma IgG, or IgG from D73, D84, or D91 plasmas, but the number of infected cells is much less in the terminal bronchioles and alveolar regions in hamsters receiving IgG from D43 plasma (b). Scale bars in magnified view denote 50 µm and those in the insets are 200 μm.

**Supp. Figure 3. Images captured in the pathological examination of the lung in hamsters treated with plasma**. Comprehensive images of the histopathology and immunohistochemistry on the lung sections of hamsters treated with control or each plasma on Day 1 post-infection are shown. Hematoxylin eosin (HE) staining of the lung sections obtained from the control plasma-(**a**), plasma D43-(**b**), plasma D73-(**c**), plasma D84-(**d**), or plasma D91-(**e**) receiving animals was done. Immunohistochemistry (IHC) for SARS-CoV-2 antigen detection of the lung sections was also conducted and shown in the right-handed side of each panel. The figures featured in Figure 4 are indicated with asterisk (*). Scale bar = 200μm

**Supp. Figure 4. Administration of IgG fraction purified from plasma D43 effectively blocks the replication and spread of SARS-CoV-2 infection in the lung tissue**. Fixed/paraffin-embedded lung tissue sections were immuno-histologically stained with anti-SARS-CoV-2-nucleoprotein polyclonal antibodies (in brown) and examined under light microscopy. Nuclei were counterstained with Mayer’s hematoxylin (in blue). Representative images of the immuno-stained lung sections of hamsters receiving IgG fraction isolated from the control-plasma (**a**) and IgG fractions from D43 (**b**), D73 (**c**), D84 (**d**), and D91 (**e**) plasmas are shown. Each inset shows a higher magnification field of the rectangle area, showing the terminal bronchioles open into the alveolar region. Note that the infected cells are observed from the terminal bronchioles into the alveolar region in animals treated with control-plasma IgG, or IgG from D73, D84, or D91 plasmas, but the number of infected cells is much less in the terminal bronchioles and alveolar regions in hamsters receiving IgG from D43 plasma (**b**). Scale bars in magnified view denote 50 µm and those in the insets are 200 μm.

**Supp. Table 1.** The amount of total human IgG in plasma transfused to hamsters. Two ml of four COVID-19 convalescent plasma and one healthy donor plasma was intraperitoneally transfused to each Syrian hamster. The concentrations of total human IgG in D43, D73, D84, and D91 plasma samples were 7.9 mg/ml, 6.2 mg/ml, 9.8 mg/ml, and 6.0 mg/ml, giving the total IgG administrated 15.8 mg, 12.4 mg, 19.6 mg, and 12.0 mg, respectively. The dosage of total human IgG per body weight in D43, D73, D84, and D91 group ranged from 191.3 – 200.0 (mg/kg), 144.2 – 148.5 (mg/kg), 219.0 – 241.4 (mg/kg), and 127.8 – 163.0 (mg/kg), respectively.

| Group | Animal # | Total IgG in 2 ml plasma (mg) | Body weight (g) | IgG per body weight (mg/kg) |
| --- | --- | --- | --- | --- |
| D43 | 24 | 15.8 | 79.3 | 199.2 |
|  | 25 | 15.8 | 79.0 | 200.0 |
|  | 26 | 15.8 | 82.6 | 191.3 |
| D73 | 27 | 12.4 | 86.0 | 144.2 |
|  | 28 | 12.4 | 83.5 | 148.5 |
|  | 29 | 12.4 | 85.1 | 145.7 |
| D84 | 30 | 19.6 | 81.2 | 241.4 |
|  | 31 | 19.6 | 89.5 | 219.0 |
|  | 32 | 19.6 | 81.5 | 240.5 |
| D91 | 33 | 12.0 | 73.6 | 163.0 |
|  | 34 | 12.0 | 93.9 | 127.8 |
|  | 35 | 12.0 | 82.7 | 145.1 |

**Supp Table 2.**
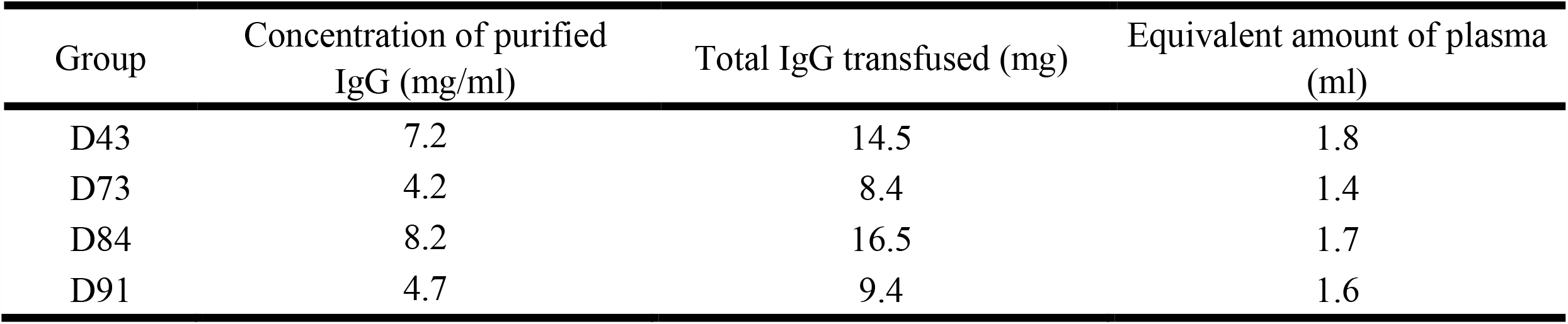
The amount of total human IgG in purified IgG transfused hamsters. Two ml of four COVID-19-convalescent plasma samples and one healthy donor plasma-derived purified IgG was intraperitoneally transfused to each Syrian hamster. The amount of total IgG transfused in D43, D73, D84, and D91 group was 14.5 mg, 8.4 mg, 16.5 mg, and 9.4 mg, which was equivalent to 1.8 ml, 1.4 ml, 1.7 ml, and 1.6 ml of the convalescent plasma, respectively.

| Group | Concentration of purified IgG (mg/ml) | Total IgG transfused (mg) | Equivalent amount of plasma (ml) |
| --- | --- | --- | --- |
| D43 | 7.2 | 14.5 | 1.8 |
| D73 | 4.2 | 8.4 | 1.4 |
| D84 | 8.2 | 16.5 | 1.7 |
| D91 | 4.7 | 9.4 | 1.6 |

## References

1 Zhu N, Zhang D, Wang W, Li X, Yang B, Song J, Zhao X, Huang B, Shi W, Lu R, Niu P, Zhan F, Ma X, Wang D, Xu W, Wu G, Gao GF, Tan W, China Novel Coronavirus Investigating and Research Team. A Novel Coronavirus from Patients with Pneumonia in China, 2019. N Engl J Med. 2020; 382, 727–733.

2 Huang C, Wang Y, Li X, Ren L, Zhao J, Hu Y, Zhang L, Fan G, Xu J, Gu X, Cheng Z, Yu T, Xia J, Wei Y, Wu W, Xie X, Yin W, Li H, Liu M, Xiao Y, Gao H, Guo L, Xie J, Wang G, Jiang R, Gao Z, Jin Q, Wang J, Cao B. Clinical features of patients infected with 2019 novel coronavirus in Wuhan, China. Lancet. 2020; 395, 497–506.

3 Mitsuya H, Kokudo N. Sustaining containment of COVID-19: global sharing for pandemic response. Glob Health Med. 2020; 2,53–55.

4 Maciosek MV, LaFrance AB, Dehmer SP, McGree DA, Flottemesch TJ, Xu Z, Solberg LI. Updated Priorities Among Effective Clinical Preventive Services. Ann Fam Med. 2017; 15, 14–22.

5 Whitney CG, Zhou F, Singleton J, Schuchat A, Centers for Disease Control and Prevention (CDC). Benefits from immunization during the vaccines for children program era - United States, 1994-2013. MMWR Morb Mortal Wkly Rep. 2014; 63, 352–355.

6 Richman DD. COVID-19 vaccines: implementation, limitations and opportunities. Glob Health Med. 2021; 3,1–5.

7 Walsh EE, Frenck RW Jr, Falsey AR, Kitchin N, Absalon J, Gurtman A, Lockhart S, Neuzil K, Mulligan MJ, Bailey R, Swanson KA, Li P, Koury K, Kalina W, Cooper D, Fontes-Garfias C, Shi PY, TüreciÖ Tompkins KR, Lyke KE, Raabe V, Dormitzer PR, Jansen KU,Şahin U, Gruber WC. Safety and Immunogenicity of Two RNA-Based Covid-19 Vaccine Candidates. N Engl J Med. 2020; 383, 2439–2450.

8 Jackson LA, Anderson EJ, Rouphael NG, Roberts PC, Makhene M, Coler RN, McCullough MP, Chappell JD, Denison MR, Stevens LJ, Pruijssers AJ, McDermott A, Flach B, Doria-Rose NA, Corbett KS, Morabito KM, O’Dell S, Schmidt SD, Swanson PA 2nd, Padilla M, Mascola JR, Neuzil KM, Bennett H, Sun W, Peters E, Makowski M, Albert J, Cross K, Buchanan W, Pikaart-Tautges R, Ledgerwood JE, Graham BS, Beigel JH, mRNA-1273 Study Group. An mRNA Vaccine against SARS-CoV-2 - Preliminary Report. N Engl J Med. 2020; 383, 1920–1931.

9 Folegatti PM, Ewer KJ, Aley PK, Angus B, Becker S, Belij-Rammerstorfer S, Bellamy D, Bibi S, Bittaye M, Clutterbuck EA, Dold C, Faust SN, Finn A, Flaxman AL, Hallis B, Heath P, Jenkin D, Lazarus R, Makinson R, Minassian AM, Pollock KM, Ramasamy M, Robinson H, Snape M, Tarrant R, Voysey M, Green C, Douglas AD, Hill AVS, Lambe T, Gilbert SC, Pollard AJ, Oxford COVID Vaccine Trial Group. Safety and immunogenicity of the ChAdOx1 nCoV-19 vaccine against SARS-CoV-2: a preliminary report of a phase 1/2, single-blind, randomised controlled trial. Lancet. 2020; 396, 467–478.

10 Sadoff J, Le Gars M, Shukarev G, Heerwegh D, Truyers C, de Groot AM, Stoop J, Tete S, Van Damme W, Leroux-Roels I, Berghmans PJ, Kimmel M, Van Damme P, de Hoon J, Smith W, Stephenson KE, De Rosa SC, Cohen KW, McElrath MJ, Cormier E, Scheper G, Barouch DH, Hendriks J, Struyf F, Douoguih M, Van Hoof J, Schuitemaker H. Interim Results of a Phase 1-2a Trial of Ad26.COV2.S Covid-19 Vaccine. N Engl J Med. 2021; 384, 1824–1835.

11 Keech C, Albert G, Cho I, Robertson A, Reed P, Neal S, Plested JS, Zhu M, Cloney-Clark S, Zhou H, Smith G, Patel N, Frieman MB, Haupt RE, Logue J, McGrath M, Weston S, Piedra PA, Desai C, Callahan K, Lewis M, Price-Abbott P, Formica N, Shinde V, Fries L, Lickliter JD, Griffin P, Wilkinson B, Glenn GM. Phase 1-2 Trial of a SARS-CoV-2 Recombinant Spike Protein Nanoparticle Vaccine. N Engl J Med. 2020; 383, 2320–2332.

12 Xia S, Duan K, Zhang Y, Zhao D, Zhang H, Xie Z, Li X, Peng C, Zhang Y, Zhang W, Yang Y, Chen W, Gao X, You W, Wang X, Wang Z, Shi Z, Wang Y, Yang X, Zhang L, Huang L, Wang Q, Lu J, Yang Y, Guo J, Zhou W, Wan X, Wu C, Wang W, Huang S, Du J, Meng Z, Pan A, Yuan Z, Shen S, Guo W, Yang X. Effect of an Inactivated Vaccine Against SARS-CoV-2 on Safety and Immunogenicity Outcomes: Interim Analysis of 2 Randomized Clinical Trials. JAMA. 2020; 324, 951–960.

13 Zhang Y, Zeng G, Pan H, Li C, Hu Y, Chu K, Han W, Chen Z, Tang R, Yin W, Chen X, Hu Y, Liu X, Jiang C, Li J, Yang M, Song Y, Wang X, Gao Q, Zhu F. Safety, tolerability, and immunogenicity of an inactivated SARS-CoV-2 vaccine in healthy adults aged 18-59 years: a randomised, double-blind, placebo-controlled, phase 1/2 clinical trial. Lancet Infect Dis. 2021; 21:181–192.

14 Ella R, Vadrevu KM, Jogdand H, Prasad S, Reddy S, Sarangi V, Ganneru B, Sapkal G, Yadav P, Abraham P, Panda S, Gupta N, Reddy P, Verma S, Kumar Rai S, Singh C, Redkar SV, Gillurkar CS, Kushwaha JS, Mohapatra S, Rao V, Guleria R, Ella K, Bhargava B. Safety and immunogenicity of an inactivated SARS-CoV-2 vaccine, BBV152: a double-blind, randomised, phase 1 trial. Lancet Infect Dis. 2021; 21:637–646.

15 Richman DD. Antiviral Drug Discovery To Address the COVID-19 Pandemic. mBio. 2020; 11, e02134–20.

16 Davies NG, Abbott S, Barnard RC, Jarvis CI, Kucharski AJ, Munday JD, Pearson CAB, Russell TW, Tully DC, Washburne AD, Wenseleers T, Gimma A, Waites W, Wong KLM, van Zandvoort K, Silverman JD, CMMID COVID-19 Working Group, COVID-19 Genomics UK (COG-UK) Consortium, Diaz-Ordaz K, Keogh R, Eggo RM, Funk S, Jit M, Atkins KE, Edmunds WJ. Estimated transmissibility and impact of SARS-CoV-2 lineage B.1.1.7 in England. Science. 2021; 372, eabg3055.

17 Xie X, Liu Y, Liu J, Zhang X, Zou J, Fontes-Garfias CR, Xia H, Swanson KA, Cutler M, Cooper D, Menachery VD, Weaver SC, Dormitzer PR, Shi PY. Neutralization of SARS-CoV-2 spike 69/70 deletion, E484K and N501Y variants by BNT162b2 vaccine-elicited sera. Nat Med. 2021; 27, 620–621.

18 Beigel JH, Tomashek KM, Dodd LE, Mehta AK, Zingman BS, Kalil AC, Hohmann E, Chu HY, Luetkemeyer A, Kline S, Lopez de Castilla D, Finberg RW, Dierberg K, Tapson V, Hsieh L, Patterson TF, Paredes R, Sweeney DA, Short WR, Touloumi G, Lye DC, Ohmagari N, Oh MD, Ruiz-Palacios GM, Benfield T, Fätkenheuer G, Kortepeter MG, Atmar RL, Creech CB, Lundgren J, Babiker AG, Pett S, Neaton JD, Burgess TH, Bonnett T, Green M, Makowski M, Osinusi A, Nayak S, Lane HC, ACTT-1 Study Group Members. Remdesivir for the Treatment of Covid-19 - Final Report. N Engl J Med. 2020; 383, 1813–1826.

19 RECOVERY Collaborative Group, Horby P, Lim WS, Emberson JR, Mafham M, Bell JL, Linsell L, Staplin N, Brightling C, Ustianowski A, Elmahi E, Prudon B, Green C, Felton T, Chadwick D, Rege K, Fegan C, Chappell LC, Faust SN, Jaki T, Jeffery K, Montgomery A, Rowan K, Juszczak E, Baillie JK, Haynes R, Landray MJ. Dexamethasone in Hospitalized Patients with Covid-19. 2021; N Engl J Med. 384, 693–704.

20 Kalil AC, Patterson TF, Mehta AK, Tomashek KM, Wolfe CR, Ghazaryan V, Marconi VC, Ruiz-Palacios GM, Hsieh L, Kline S, Tapson V, Iovine NM, Jain MK, Sweeney DA, El Sahly HM, Branche AR, Regalado Pineda J, Lye DC, Sandkovsky U, Luetkemeyer AF, Cohen SH, Finberg RW, Jackson PEH, Taiwo B, Paules CI, Arguinchona H, Erdmann N, Ahuja N, Frank M, Oh MD, Kim ES, Tan SY, Mularski RA, Nielsen H, Ponce PO, Taylor BS, Larson L, Rouphael NG, Saklawi Y, Cantos VD, Ko ER, Engemann JJ, Amin AN, Watanabe M, Billings J, Elie MC, Davey RT, Burgess TH, Ferreira J, Green M, Makowski M, Cardoso A, de Bono S, Bonnett T, Proschan M, Deye GA, Dempsey W, Nayak SU, Dodd LE, Beigel JH, ACTT-2 Study Group Members. Baricitinib plus Remdesivir for Hospitalized Adults with Covid-19. N Engl J Med. 2021; 384, 795–807.

21 Mehta P, McAuley DF, Brown M, Sanchez E, Tattersall RS, Manson JJ, HLH Across Speciality Collaboration, UK. COVID-19: consider cytokine storm syndromes and immunosuppression. Lancet. 2020; 395, 1033–1034.

22 Stone JH, Frigault MJ, Serling-Boyd NJ, Fernandes AD, Harvey L, Foulkes AS, Horick NK, Healy BC, Shah R, Bensaci AM, Woolley AE, Nikiforow S, Lin N, Sagar M, Schrager H, Huckins DS, Axelrod M, Pincus MD, Fleisher J, Sacks CA, Dougan M, North CM, Halvorsen YD, Thurber TK, Dagher Z, Scherer A, Wallwork RS, Kim AY, Schoenfeld S, Sen P, Neilan TG, Perugino CA, Unizony SH, Collier DS, Matza MA, Yinh JM, Bowman KA, Meyerowitz E, Zafar A, Drobni ZD, Bolster MB, Kohler M, D’Silva KM, Dau J, Lockwood MM, Cubbison C, Weber BN, Mansour MK, BACC Bay Tocilizumab Trial Investigators. Efficacy of Tocilizumab in Patients Hospitalized with Covid-19. N Engl J Med. 2020; 383, 2333–2344.

23 Wang Y, Zhang D, Du G, Du R, Zhao J, Jin Y, Fu S, Gao L, Cheng Z, Lu Q, Hu Y, Luo G, Wang K, Lu Y, Li H, Wang S, Ruan S, Yang C, Mei C, Wang Y, Ding D, Wu F, Tang X, Ye X, Ye Y, Liu B, Yang J, Yin W, Wang A, Fan G, Zhou F, Liu Z, Gu X, Xu J, Shang L, Zhang Y, Cao L, Guo T, Wan Y, Qin H, Jiang Y, Jaki T, Hayden FG, Horby PW, Cao B, Wang C. Remdesivir in adults with severe COVID-19: a randomised, double-blind, placebo-controlled, multicentre trial. Lancet. 2020; 395, 1569–1578.

24 Carter PJ, Lazar GA. Next generation antibody drugs: pursuit of the ‘high-hanging fruit’. Nat Rev Drug Discov. 2018; 17, 197–223.

25 Piechotta V, Iannizzi C, Chai KL, Valk SJ, Kimber C, Dorando E, Monsef I, Wood EMLamikanra AA, Roberts DJ, McQuilten Z, So-Osman C, Estcourt LJ, Skoetz N. Convalescent plasma or hyperimmune immunoglobulin for people with COVID-19: a living systematic review. Cochrane Database Syst Rev. 2021; 5, CD013600.

26 Salazar E, Christensen PA, Graviss EA, Nguyen DT, Castillo B, Chen J, Lopez BV, Eagar TN, Yi X, Zhao P, Rogers J, Shehabeldin A, Joseph D, Masud F, Leveque C, Olsen RJ, Bernard DW, Gollihar J, Musser JM. Significantly Decreased Mortality in a Large Cohort of Coronavirus Disease 2019 (COVID-19) Patients Transfused Early with Convalescent Plasma Containing High-Titer Anti–Severe Acute Respiratory Syndrome Coronavirus 2 (SARS-CoV-2) Spike Protein IgG. Am J Pathol. 2021; 191, 90–107.

27 Libster R, Pérez Marc G, Wappner D, Coviello S, Bianchi A, Braem V, Esteban I, Caballero MT, Wood C, Berrueta M, Rondan A, Lescano G, Cruz P, Ritou Y, Fernández Viña V,Álvarez Paggi D, Esperante S, Ferreti A, Ofman G, CigandaÁ Rodriguez R, Lantos J, Valentini R, Itcovici N, Hintze A, Oyarvide ML, Etchegaray C, Neira A, Name I, Alfonso J, López Castelo R, Caruso G, Rapelius S, Alvez F, Etchenique F, Dimase F, Alvarez D, Aranda SS, Sánchez Yanotti C, De Luca J, Jares Baglivo S, Laudanno S, Nowogrodzki F, Larrea R, Silveyra MLeberzstein G, Debonis A, Molinos J, González M, Perez E, Kreplak N, Pastor Argüello S, Gibbons L, Althabe F, Bergel E, Polack FP, Fundación INFANT–COVID-19 Group. Early High-Titer Plasma Therapy to Prevent Severe Covid-19 in Older Adults. N Engl J Med. 2021; 384, 610–618.

28 Imai M, Iwatsuki-Horimoto K, Hatta MLoeber S, Halfmann PJ, Nakajima N, Watanabe T, Ujie M, Takahashi K, Ito M, Yamada S, Fan S, Chiba S, Kuroda M, Guan L, Takada K, Armbrust T, Balogh A, Furusawa Y, Okuda M, Ueki H, Yasuhara A, Sakai-Tagawa Y, Lopes TJS, Kiso M, Yamayoshi S, Kinoshita N, Ohmagari N, Hattori SI, Takeda M, Mitsuya H, Krammer F, Suzuki T, Kawaoka Y. Syrian hamsters as a small animal model for SARS-CoV-2 infection and countermeasure development. Proc Natl Acad Sci U S A. 2020; 117, 16587–16595.

29 Hattori SI, Higashi-Kuwata N, Hayashi H, Allu SR, Raghavaiah J, Bulut H, Das D, Anson BJ, Lendy EK, Takamatsu Y, Takamune N, Kishimoto N, Murayama K, Hasegawa K, Li M, Davis DA, Kodama EN, Yarchoan R, Wlodawer A, Misumi S, Mesecar AD, Ghosh AK, Mitsuya H. A small molecule compound with an indole moiety inhibits the main protease of SARS-CoV-2 and blocks virus replication. Nat Commun. 2021; 12, 668.

30 Maeda K, Higashi-Kuwata N, Kinoshita N, Kutsuna S, Tsuchiya K, Hattori SI, Matsuda K, Takamatsu Y, Gatanaga H, Oka S, Sugyama H, Ohmagari N, Mitsuya H. Neutralization of SARS-CoV-2 with IgG from COVID-19-convalescent plasma. Sci Rep. 2021; 11, 5563.

31 Hattori SI, Higashi-Kuwata N, Raghavaiah J, Das D, Bulut H, Davis DA, Takamatsu Y, Matsuda K, Takamune N, Kishimoto N, Okamura T, Misumi S, Yarchoan R, Maeda K, Ghosh AK, Mitsuya H. GRL-0920, an Indole Chloropyridinyl Ester, Completely Blocks SARS-CoV-2 Infection. mBio. 2020; 11, e01833–20.

32 Draper NR, Smith H. Applied regression analysis. John Wiley & Sons, New York, 1966.

33 Grifoni A, Weiskopf D, Ramirez SI, Mateus J, Dan JM, Moderbacher CR, Rawlings SA, Sutherland A, Premkumar L, Jadi RS, Marrama D, de Silva AM, Frazier A, Carlin AF, Greenbaum JA, Peters B, Krammer F, Smith DM, Crotty S, Sette A. Targets of T Cell Responses to SARS-CoV-2 Coronavirus in Humans with COVID-19 Disease and Unexposed Individuals. Cell. 2020; 181, 1489–1501.

34 Dan JM, Mateus J, Kato Y, Hastie KM, Yu ED, Faliti CE, Grifoni A, Ramirez SI, Haupt S, Frazier A, Nakao C, Rayaprolu V, Rawlings SA, Peters B, Krammer F, Simon V, Saphire EO, Smith DM, Weiskopf D, Sette A, Crotty S. Immunological memory to SARS-CoV-2 assessed for up to 8 months after infection. Science. 2021; 371, eabf4063.

35 Kristiansen PA, Page M, Bernasconi V, Mattiuzzo G, Dull P, Makar K, Plotkin S, Knezevic I. WHO International Standard for anti-SARS-CoV-2 immunoglobulin. Lancet. 2021; 397, 1347–1348.

36 Noy-Porat T, Makdasi E, Alcalay R, Mechaly A, Levy Y, Bercovich-Kinori A, Zauberman A, Tamir H, Yahalom-Ronen Y, Israeli M, Epstein E, Achdout H, Melamed S, Chitlaru T, Weiss S, Peretz E, Rosen O, Paran N, Yitzhaki S, Shapira SC, Israely T, Mazor O, Rosenfeld R. A panel of human neutralizing mAbs targeting SARS-CoV-2 spike at multiple epitopes. Nat Commun. 2020; 11, 4303

37 Rodda LB, Netland J, Shehata L, Pruner KB, Morawski PA, Thouvenel CD, Takehara KK, Eggenberger J, Hemann EA, Waterman HR, Fahning ML, Chen Y, Hale M, Rathe J, Stokes C, Wrenn S, Fiala B, Carter L, Hamerman JA, King NP, Gale M Jr, Campbell DJ, Rawlings DJ, Pepper M. Functional SARS-CoV-2-Specific Immune Memory Persists after Mild COVID-19. Cell. 2021; 184, 169–183.

38 Baric RS. Emergence of a Highly Fit SARS-CoV-2 Variant. N Engl J Med. 2020; 383, 2684–2686.

39 Frampton D, Rampling T, Cross A, Bailey H, Heaney J, Byott M, Scott R, Sconza R, Price J, Margaritis M, Bergstrom M, Spyer MJ, Miralhes PB, Grant P, Kirk S, Valerio C, Mangera Z, Prabhahar T, Moreno-Cuesta J, Arulkumaran N, Singer M, Shin GY, Sanchez E, Paraskevopoulou SM, Pillay D, McKendry RA, Mirfenderesky M, Houlihan CF, Nastouli E. Genomic characteristics and clinical effect of the emergent SARS-CoV-2 B.1.1.7 lineage in London, UK: a whole-genome sequencing and hospital-based cohort study. Lancet Infect Dis. 2021; S1473-3099(21)00170-5. doi: 10.1016/S1473-3099(21)00170-5. Online ahead of print.

40 Tegally H, Wilkinson E, Giovanetti M, Iranzadeh A, Fonseca V, Giandhari J, Doolabh D, Pillay S, San EJ, Msomi N, Mlisana K, von Gottberg A, Walaza S, Allam M, Ismail A, Mohale T, Glass AJ, Engelbrecht S, Van Zyl G, Preiser W, Petruccione F, Sigal A, Hardie D, Marais G, Hsiao NY, Korsman S, Davies MA, Tyers L, Mudau I, York D, Maslo C, Goedhals D, Abrahams S, Laguda-Akingba O, Alisoltani-Dehkordi A, Godzik A, Wibmer CK, Sewell BT, Lourenço J, Alcantara LCJ, Kosakovsky Pond SL, Weaver S, Martin D, Lessells RJ, Bhiman JN, Williamson C, de Oliveira T. Detection of a SARS-CoV-2 variant of concern in South Africa. Nature. 2021; 592, 438–443.

41 Faria NR, Mellan TA, Whittaker C, Claro IM, Candido DDS, Mishra S, Crispim MAE, Sales FCS, Hawryluk I, McCrone JT, Hulswit RJG, Franco LAM, Ramundo MS, de Jesus JG, Andrade PS, Coletti TM, Ferreira GM, Silva CAM, Manuli ER, Pereira RHM, Peixoto PS, Kraemer MUG, Gaburo N Jr, Camilo CDC, Hoeltgebaum H, Souza WM, Rocha EC, de Souza LM, de Pinho MC, Araujo LJT, Malta FSV, de Lima AB, Silva JDP, Zauli DAG, Ferreira ACS, Schnekenberg RP, Laydon DJ, Walker PGT, Schlüter HM, Dos Santos ALP, Vidal MS, Del Caro VS, Filho RMF, Dos Santos HM, Aguiar RS, Proença-Modena JL, Nelson B, Hay JA, Monod M, Miscouridou X, Coupland H, Sonabend R, Vollmer M, Gandy A, Prete CA Jr, Nascimento VH, Suchard MA, Bowden TA, Pond SLK, Wu CH, Ratmann O, Ferguson NM, Dye C, Loman NJ, Lemey P, Rambaut A, Fraiji NA, Carvalho MDPSS, Pybus OG, Flaxman S, Bhatt S, Sabino EC. Genomics and epidemiology of the P.1 SARS-CoV-2 lineage in Manaus, Brazil. Science. 2021; 372, 815–821.

42 Shen X, Tang H, Pajon R, Smith G, Glenn GM, Shi W, Korber B, Montefiori DC. Neutralization of SARS-CoV-2 Variants B.1.429 and B.1.351. N Engl J Med. 2021; NEJMc2103740. doi: 10.1056/NEJMc2103740. Online ahead of print.

43 Zhou B, Thao TTN, Hoffmann D, Taddeo A, Ebert N, Labroussaa F, Pohlmann A, King J, Steiner S, Kelly JN, Portmann J, Halwe NJ, Ulrich L, Trüeb BS, Fan X, Hoffmann B, Wang L, Thomann L, Lin X, Stalder H, Pozzi B, de Brot S, Jiang N, Cui D, Hossain J, Wilson MM, Keller MW, Stark TJ, Barnes JR, Dijkman R, Jores J, Benarafa C, Wentworth DE, Thiel V, Beer M. SARS-CoV-2 spike D614G change enhances replication and transmission. Nature. 2021; 592, 122–127.

